# Herd Immunity Alters the Conditions for Performing Dose Schedule Comparisons: An Individual-based Model of Pneumococcal Carriage

**DOI:** 10.1101/393108

**Authors:** Alan Yang, Francisco Cai, Marc Lipsitch

## BACKGROUND

Mass vaccination of infants and toddlers with pneumococcal conjugate vaccines (PCVs) has led to large declines in pneumococcal disease in countries around the world [1]. These vaccines currently contain 10 or 13 different capsular polysaccharides from *Streptococcus pneumoniae*, each conjugated to a protein carrier. In *S. pneumoniae*, the chemical structure of the capsular polysaccharide determines the serotype, defined by the specificity of antibody responses against these capsules. PCVs induce immunity that directly protects recipients against vaccine serotypes (VT), which can cause invasive and mucosal disease [2]. It is clear that the immunity induced by PCVs also partially protects recipients against nasopharyngeal carriage of VT pneumococci [3]. This is important because the nasopharyngeal pneumococcal population is the source of transmission to other hosts, so the reduced carriage has led to herd immunity that has reduced VT disease in other age groups in many populations in which infants and/or toddlers have received PCVs [4, 5].

Dosing schedules adopted in national programs to date vary, with either 2 or 3 doses in infancy, with or without a booster around 12 months of age (schedules are referred to by the number of doses in infancy + the number of boosters, e.g. 2+1 or 3+0). The high market cost of PCVs has created intense interest in the possibility of reducing the total number of doses administered [6]. This interest was recently bolstered by a randomized, controlled trial showing good immunogenicity of a 1+1 schedule compared to a 2+1 schedule in the UK [7, 8]. However, to our knowledge, no randomized trial testing the impact of reduced dosing schedules on an efficacy outcome, such as prevalence of VT pneumococcal carriage, has been carried out in a country that has already implemented routine vaccination. Instead, such trials are being carried out in populations that have not yet introduced routine pneumococcal vaccination [9]. If these trials were to be conducted in countries with established vaccination programs, they would be challenged by the presence of herd immunity in the background population. We undertook simulation modeling of such trials in newborns in both the presence and absence of background herd immunity to understand the effects of background herd immunity on the ability of a vaccine trial to reveal differences in efficacy against vaccine-type carriage between its arms. We found that the differences between arms in VT pneumococcal colonization in such a trial conducted in a background of herd immunity would be very small, and the resulting sample size requirement would be impracticably large. These findings support the wisdom of locating such trials with carriage outcomes in countries that have not yet introduced routine PCV programs.

## METHODS

To simulate *S. pneumoniae* transmission dynamics, we used a discrete-time individual-based model based on a previous model developed by Cobey *et al.* [10]. As in Cobey *et al.*’s model, our model includes both serotype-specific and non-serotype-specific immunity and reproduces observed serotype diversity. The calculation of the force of colonization and duration of colonization also follow Cobey *et al*. In particular, the force of colonization for each individual is calculated every day with age-assortative mixing in transmission. Furthermore, individuals can be colonized by multiple strains of the same serotype at the same time, but the model is based on a “neutral null model” that avoids artificial predictions of serotype coexistence [11].

Our model differed from that of Cobey *et al*. in three ways. First, some versions of Cobey *et al.*’s model take into account a range of demographic events such as partnering, reproduction, and departure from the household of origin. By contrast, our model does not include these events because they are not central to the question we are interested in, which is the effect of herd immunity on the results of vaccine trials. Second, our procedure for fitting the transmission rate (β) deviated from that of Cobey *et al*.’s study. Cobey *et al*. estimated the total prevalence of pneumococcal carriage by averaging the annual samples in the final 10 years of their simulations, whereas we average over the final 46 years of our simulations to obtain more precise estimates. The transmission parameters were fit using 2001 data from the SPARC project, which sampled children aged 7 or younger in Massachusetts, USA and reported a 28% carriage prevalence [12]. We fit to serotype-specific prevalences using an algorithm previously described [13].

The third difference was that we implemented a vaccine trial. In real-world vaccine trials, the effect of the trial on background transmission is minimized by having a trial that is small relative to the general population. In our simulations, participants in our vaccine trials were modeled as individuals who could receive colonizations from individuals in the general population but could not themselves transmit colonizations to any other individuals. This design allows us to introduce a large number of vaccine trial participants to the simulation—and thus obtain stable and precise estimates of the prevalence of VT colonization in the different arms—without interfering with the level of colonization in the background population. These prevalence estimates were then used as the “true” population means in our sample size calculations.

We simulated a vaccine trial with three arms receiving the 13-valent pneumococcal vaccine (PCV13) on different schedules—3+1, 2+1, and 1+1—where the booster was always given at 12 months of age and the primary sequence was at 2, 4, and 6 months of age, with the reduced schedules omitting the 6 month, or the 4 and 6 month doses respectively. These trials took place in one of two settings: first, in a setting where the 7-valent PCV (PCV7) had been in routine use for some years and had recently been replaced by PCV13 (see details below), and second, in a PCV-naïve population. Our simulation consisted of three sequential phases:

1. **Demographic Phase (50 years)**: There is no transmission and the goal is the reach an equilibrium age distribution in the background population mimicking that in the US population. Vaccine trial has not been initiated yet. Individuals’ ages and colonizations are recorded every year.
2. **Epidemiologic Phase (50 years)**: At the beginning of this phase, we introduce the disease to the population by randomly seeding each individual in the background population with colonizations and allowing transmission to take place and equilibrate. Initially, as a temporary measure, individuals are artificially given some pre-existing immunity to prevent a computationally intensive epidemic that would have otherwise occurred had the population been fully susceptible. If simulating in the presence of routine PCV vaccination, we begin introducing PCV7 to newborns in the background population 40 years into this phase. Then, 44 years into the phase, we replace PCV7 with PCV13 in the background population. This timeline was chosen to match pneumococcal vaccination in the UK, where PCV7 was administered for 4 years before switching to PCV13 [14]. In this phase of the simulation, the vaccine trial has not started yet. Individuals’ ages and colonizations are recorded every year.
3. **Vaccine Trial Phase (5 years)**: We begin the vaccine trial by introducing newborn participants to the simulation who are able to receive colonizations but unable to transmit them. If we are simulating the vaccine trial in the presence of herd immunity from PCV, the background population continues to receive PCV13. Individuals’ ages and colonizations are recorded more frequently—every month.

We simulated 50,000 individuals in the background population (our populations size stayed constant, as in Cobey *et al*.) and 10,000 individuals in each of the 3 arms of the trial. Vaccines were assumed to confer varying degrees of protection with increasing numbers of doses, with the increased efficacy on subsequent doses taking effect immediately on receipt of the next dose. Vaccine efficacy after two or more doses (for 2+1 and 3+1 PCV13 schedules) was estimated using VT prevalence from Dagan *et al*. [15] and calculated as one minus the prevalence odds ratio as described in Rinta-Kokko *et al.* (**Table 1**) [16]. Vaccine efficacy after one dose (for all three PCV13 schedules) was estimated using a similar approach with data from Ota *et al*. [17] To our knowledge, no data was available for the efficacy of the 1+1 schedule after the booster dose. Because initial analyses (data not shown) had found very large sample sizes for comparative trials, we made the assumption that would yield the greatest difference between arms and thus the smallest sample size: that efficacy remained unchanged after the booster dose in the 1+1 schedule. Other equally plausible assumptions would have resulted in greater protection of this trial group and thus would have required even larger sample sizes to see a difference with other arms. PCV7 efficacy was assumed to follow the same schedule as PCV13, but limited to the relevant serotypes.

**Table 1.**
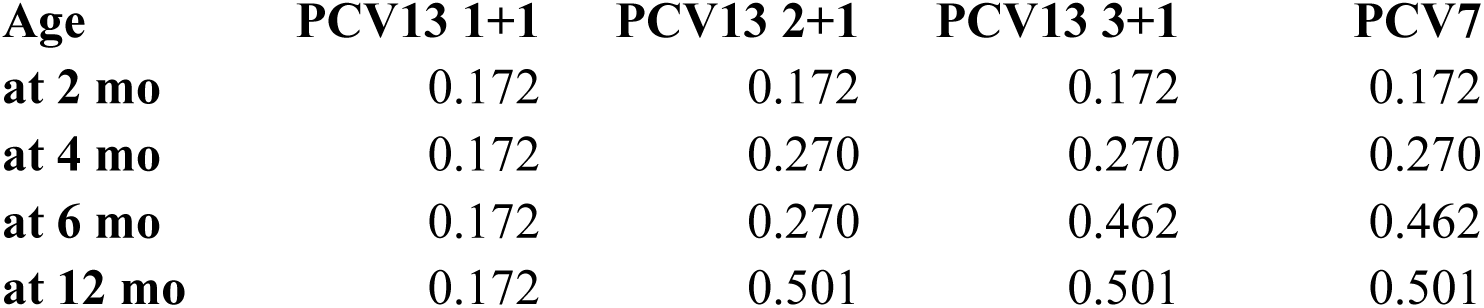
Vaccine Schedule and Efficacies as Modeled in This Simulation.

We ran each simulation 10 times and averaged the results of these iterations to obtain fairly smooth prevalence estimates. The simulation was coded in C++11 and run using the software XCode (Version 7.2.1). Data analysis was done in Jupyter Notebook (Version 4.0.6).

We calculated the sample sizes needed to achieve a certain power in distinguishing two trial arms by taking the effect size to be the maximum difference in VT prevalence between the two trials during the course of the entire trial. We then determined the relationship between power and sample size by interpreting the VT prevalence of each trial as a proportion. The specific formula we used to calculate this relationship follows Cohen [18].

In addition to sample size, we calculated what estimate or relative efficacy would emerge at different timepoints in the trial. Relative efficacy was defined as either one minus the risk ratio, or one minus the odds ratio, of VT carriage prevalence between arms of the trial, with the 1+1 arm as the reference group and the higher-dose groups as the “interventions.”

## RESULTS

**In a PCV-naïve population, the difference in VT pneumococcal carriage prevalence between trial arms was less than 7% and varied with sampling time.** The largest prevalence difference between the 3+1 and 1+1 trial arms, 6.4%, was observed 34 months after the start of the vaccine trial. Meanwhile, the largest difference between the 3+1 and the 2+1 trial arms was 2.3%, occurring 12 months into the vaccine trial. **(Figure 1, a-c)**

**Figure 1:**
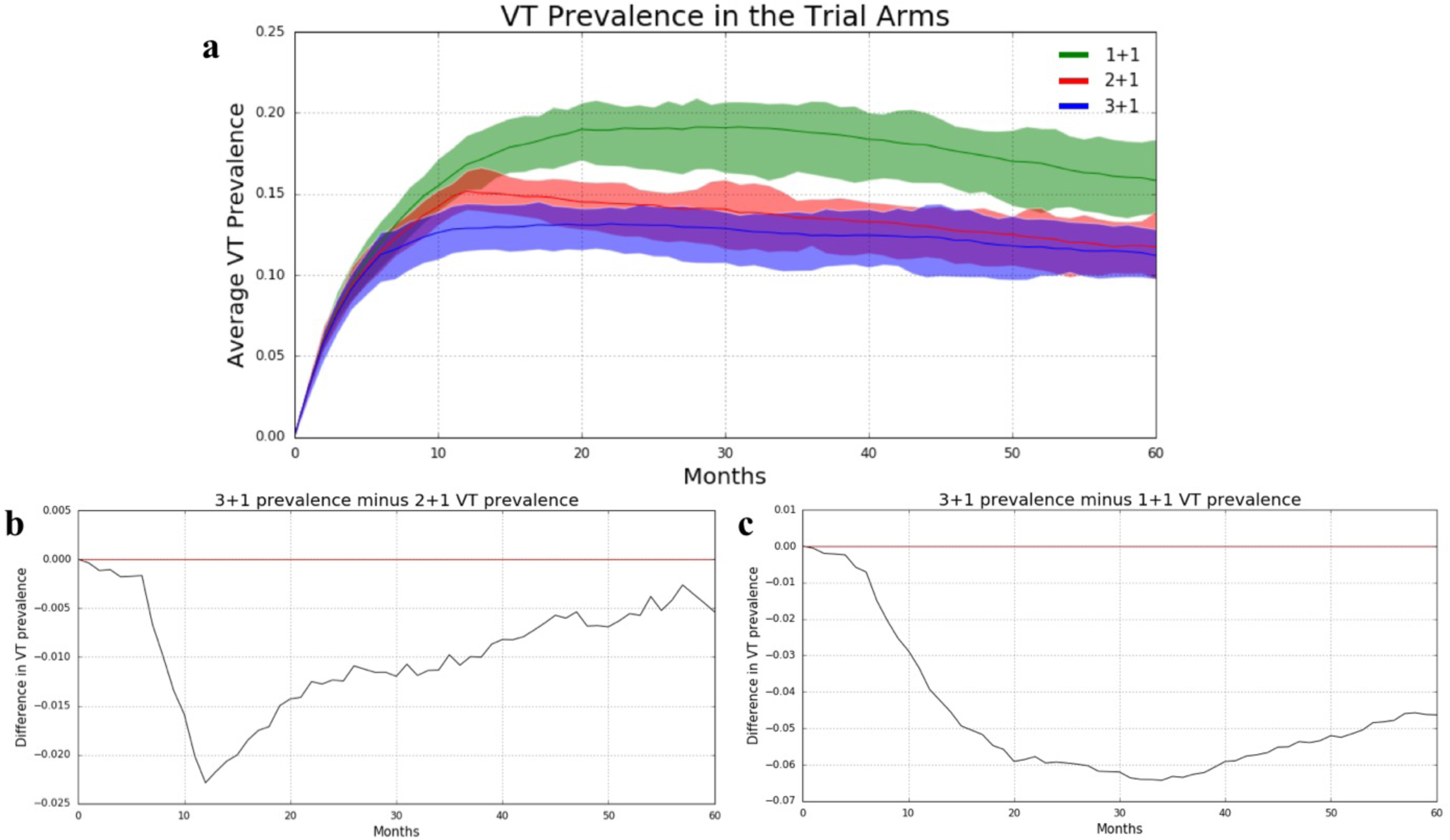
Naïve Population. (a) VT carriage prevalence in individual trial arms. Lines represent average VT prevalence across 10 simulations while shading is bounded by the maximum and minimum VT prevalences obtained across the 10 simulation at any given point in time. (b) The difference in VT prevalence between the 3+1 and 2+1 trial arms (averaged across 10 simulations) graphed over time. (c) The difference in VT prevalence between the 3+1 and 1+1 trial arms (averaged across 10 simulations) graphed over time. The zero difference line is shown in red.

**In a population already receiving routine PCV administration, VT pneumococcal prevalence is nearly indistinguishable between trial arms.** The overall prevalences in each trial arm are much lower than in a PCV-naïve population, and the differences between the trial arms are very small. Indeed, the largest difference between the 3+1 and 1+1 trial arms was 1.0%, which occurred 24 months into the vaccine trial. The largest difference between the 3+1 and 2+1 trial arms was only 0.5% and occurred 12 months into the vaccine trial. **(Figure 2, a-c)**

**Figure 2:**
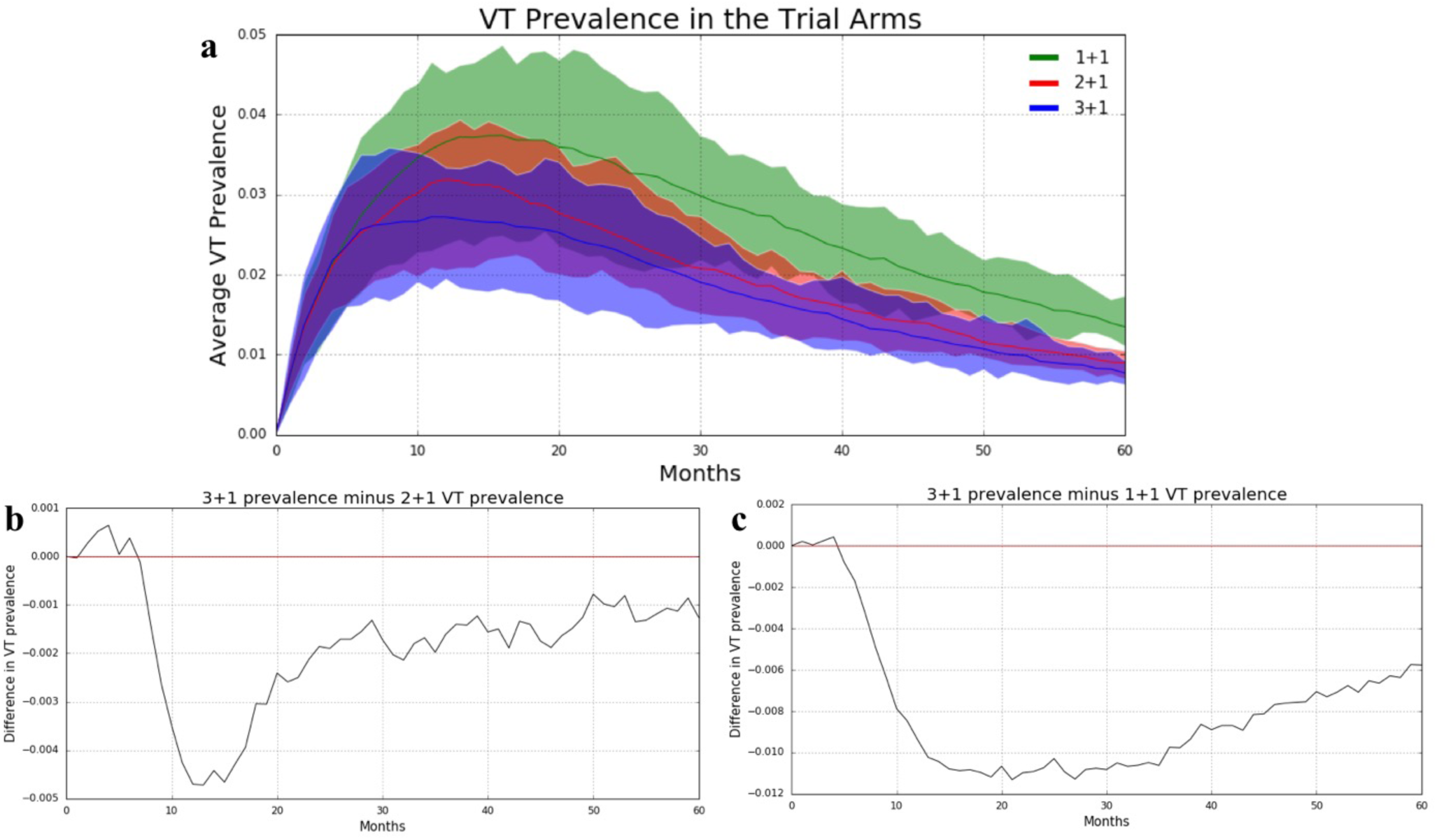
3+1 Vaccinated Population. (a) VT carriage prevalence in individual trial arms. Lines represent average VT prevalence across 10 simulations while shading is bounded by the maximum and minimum VT prevalences obtained across the 10 simulation at any given point in time. (b) The difference in VT prevalence between the 3+1 and 2+1 trial arms (averaged across 10 simulations) graphed over time. (c) The difference in VT prevalence between the 3+1 and 1+1 trial arms (averaged across 10 simulations) graphed over time. The zero difference line is shown in red. Note the scale of the y-axis.

**Relative efficacy estimation.** Vaccine efficacy (VE) against carriage is often calculated as VE = 1 – (prevalence of VT carriage in vaccinees)/(prevalence of VT carriage in controls), or alternatively as 1 – (prevalence odds of VT carriage in vaccinees)/(prevalence odds of VT carriage in controls) [1]. Figure 3 compares VE by each of these two measures over time, during the vaccine trial. Notably, vaccine efficacy varies with time since vaccination, due to the dynamic nature of pneumococcal colonization [13]. Also interestingly, the vaccine efficacy measured post-vaccination is generally lower in a trial in a vaccine-naïve community than one in a community already using PCV13. Moreover, this difference in VE grows larger as time goes on (**Figure 3**)

**Figure 3:**
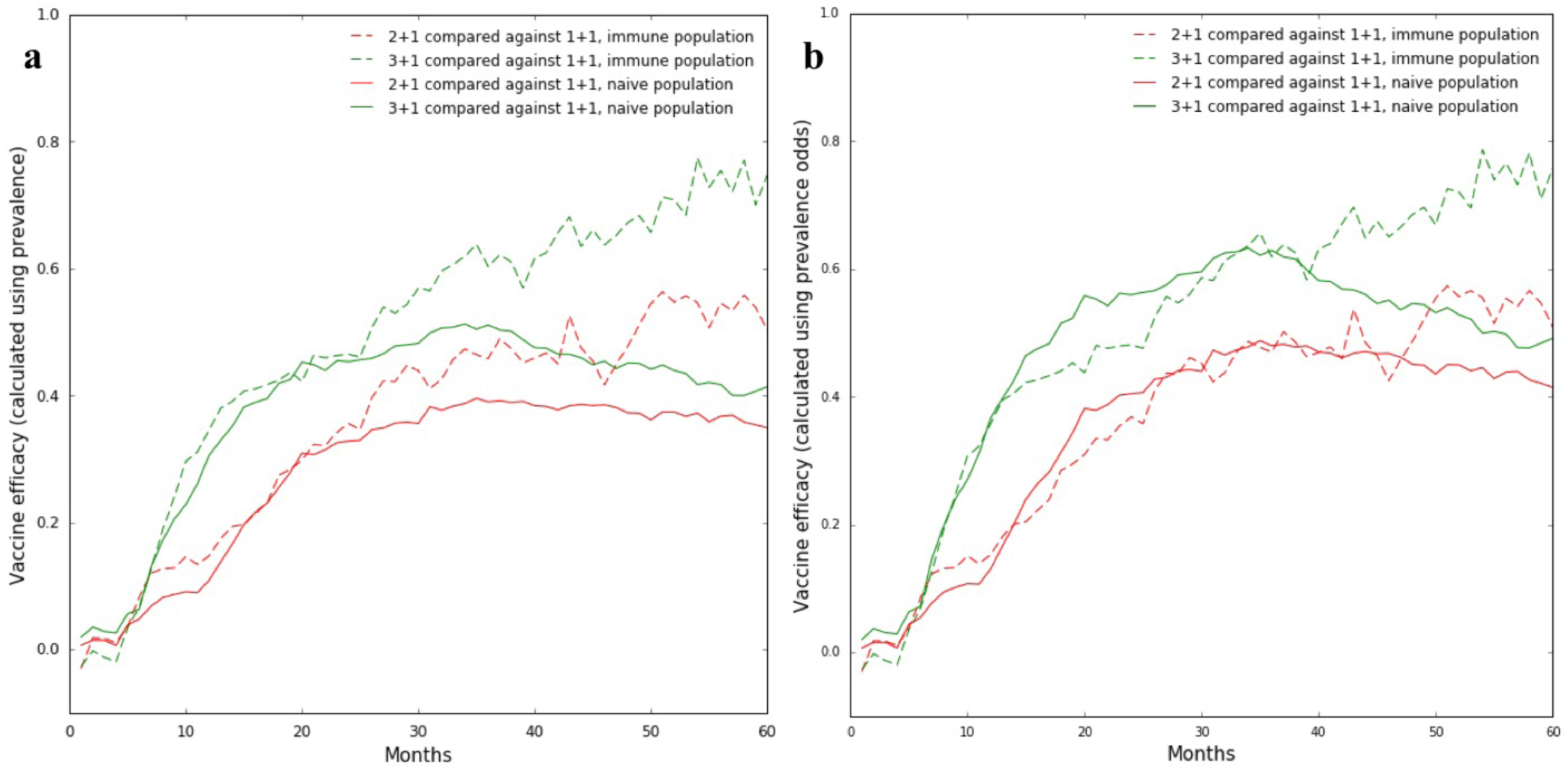
Relative Efficacies During the Vaccine Trial. (a) VE calculated using prevalences. (b) VE calculated using prevalence odds.

**Much larger sample sizes—by an order of magnitude—are required for a vaccine trial conducted in a population receiving routine PCV administration as compared to in PCV-naïve population**.

In a PCV-naïve population the sample size needed to distinguish the 3+1 arm from the 1+1 arm with a power of 80% is roughly 290 (*α* = 0.05); distinguishing the 3+1 arm from the 2+1 arm with a power of 80% requires a sample size of 2095 (again, *α* = 0.05). By contrast, in a population already receiving routine PCV administration, the sample size needed to distinguish the 3+1 arm from the 1+1 arm with a power of 80% is about 2070; distinguishing the 3+1 arm from the 1+1 arm here with a power of 80% requires a sample size of nearly 11620. **(Figure 4)**

**Figure 4:**
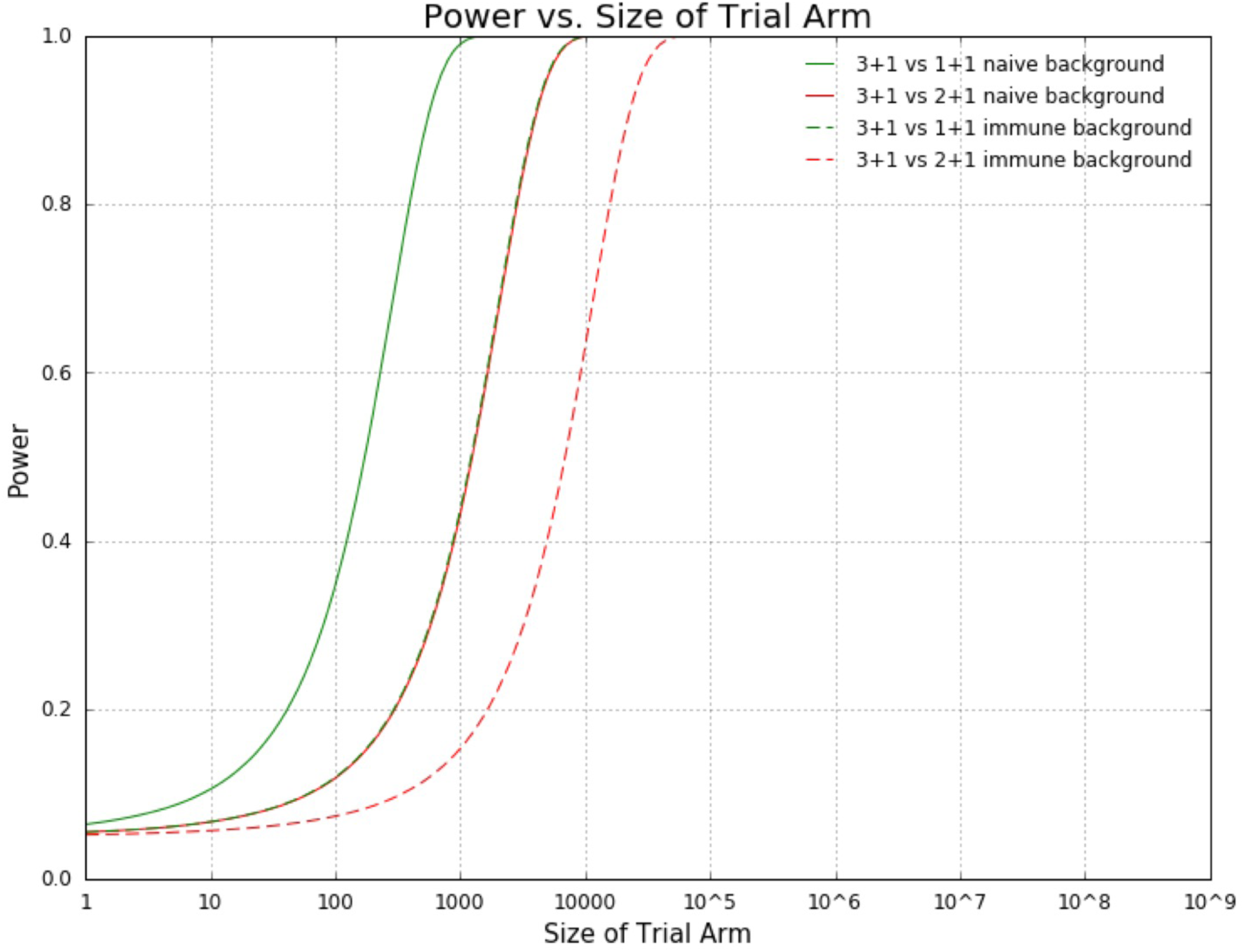
Power vs. Trial arm size.

## DISCUSSION

This simulation study showed that in the presence of significant herd immunity conferred by widespread uptake of PCVs, a clinical trial to estimate the effects of reduced dosing schedules on VT pneumococcal carriage would require impracticably large sample sizes. This occurs both because the level of VT carriage is reduced in the target age group and by the herd immunity effects.

These findings confirm the wisdom of the decision to test immunogenicity of reduced dosing schedules, but not efficacy against carriage, in countries that have already adopted PCV vaccination as a national policy [8], while conducting head-to-head randomized comparisons of dosing schedules only in countries which have not yet introduced a policy of PCV use, such as Viet Nam [3, 19].

The result that it would be difficult to detect a small reduction in immune protection in a context of rare VT colonization was perhaps predictable by simple intuitive reasoning. However, previous work [13] has shown that pneumococcal vaccine trials with a colonization prevalence endpoint have several particularities that differ from those of more traditional vaccine trials using incidence of an acute infection as an endpoint [20, 21, 22, 23, 24]. In particular, these studies have shown that several factors complicate the interpretation of such studies:

- the competition between different serotypes to colonize [25, 26]
- the ability of most assays to detect only one serotype among those colonized with one or more serotypes [27]
- the nonlinear relationship between incidence rate reduction (the biological parameter assumed to be relatively consistent for the same vaccine in different settings) and prevalence reduction (which is measured in such trials) [16]
- the fact that as children are receiving vaccine doses they are also acquiring immunity in both serotype-specific and nonspecific fashions to colonization. [24]

It is thus risky to rely on intuition for how vaccine trials would work in various settings; hence, the impetus to perform the studies described here. Indeed, one part of our results – the finding that vaccine efficacy as measured by reduction in either risk or odds of carriage is higher in the presence of herd immunity – is counter to the usual pattern in vaccine trials using incidence as an endpoint, wherein reduced incidence in all trial arms may occur due to herd immunity, but the relative reduction in incidence due to vaccination (or more vaccination) is assumed to be constant across settings [28]. The strong time-dependence of measured relative efficacy of various dosing schedule was also a striking finding that has not been considered in most previous studies; this finding again is a result of the use of prevalence rather than incidence as an endpoint, and of the complex dynamics of age-varying force of infection and developing immunity in the early years of life via natural colonization [24]. In general, simulations of clinical trials for vaccines or other infectious disease prevention measures can enhance intuition and improve study design [29].

Like any model, the simulation model here has several limitations. While it attempts to incorporate the major determinants of both serotype-specific (antibody-based, acquisition-reducing) and serotype-transcending (Th17-based, duration-reducing) immunity to carriage of pneumococci, as well as competition between different strains to colonize the same host, the design of the model required a number of simplifying assumptions, in particular that these were the only relevant forms of constraint on acquisition of pneumococcal carriage. We did not incorporate household structure in these simulations, because earlier work [10] showed that qualitative patterns were not strongly affected by these details. Carriage prevalence by serotype was calibrated to data from Massachusetts, USA [30], and the details of carriage prevalence overall can vary considerably across populations, though the leading serotypes (prior to vaccine introduction) are remarkably consistent. The assumed reductions in carriage incidence due to various dosing schedules were point estimates taken from a single study, and these are subject both to statistical uncertainty and possible variation across settings [10]. Despite these limitations, we believe that the large magnitude of the simulated difference between vaccine-experienced and vaccine-naïve settings shows robustly that such trials would be almost certainly underpowered if conducted in populations that have already introduced mass pneumococcal conjugate vaccination. Moreover, particularly in light of recent studies of a different vaccine and its likely performance in clinical trials, the finding of time-dependent variation in efficacy appears to be a robust result [24]. Finally, we calculated sample sizes for the point of largest prevalence difference in the simulations, for a superiority study. Sample sizes for a noninferiority study may differ depending on the margin of noninferiority chosen.

Further work is needed to understand the long-term implications of such altered vaccine schedules as, over time, persons who received reduced schedules as infants constitute a growing proportion of the population.

## CONCLUSIONS

This study demonstrates computationally that the presence of background herd immunity challenges the comparison of PCV dose schedules in a clinical trial. In particular, immune populations require impractical samples sizes that are an order of magnitude larger than those for vaccine-naïve populations in order to distinguish between the carriage prevalences of the arms of a PCV dosage-comparison trial. This result holds true, moreover, despite the fact that VE estimates are generally higher in immune populations than in vaccine-naïve populations.

Not only are the results of a PCV dose schedule comparison trial affected by background herd immunity, but they are also time-dependent. Specifically, the relative efficacy of different dosing schedules varies strongly with time, with maximal prevalence differences attained 1-3 years into the trial.

By highlighting the context- and time-dependence of efficacy estimates in PCV dose schedule comparison trials, these findings underscore some underappreciated aspects of these trials and support the wisdom of comparing differences in carriage between individuals receiving different dosages of PCV only in vaccine-naïve populations.

## LIST OF ABBREVIATIONS

PCV: Pneumococcal conjugate vaccine

VT: Vaccine type

PCV7: 7-valent pneumococcal conjugate vaccine

PCV13: 13-valent pneumococcal conjugate vaccine

VE: vaccine efficacy

## DECLARATIONS

### Ethics approval and consent to participate

Not applicable

### Consent for publication

Not applicable

### Availability of data and material

The datasets generated and analyzed during the current study are available from the corresponding author upon request.

### Authors’ contributions

AY and FC helped modified the simulation code first written by Cobey *et al*. [10]. AY, FC, and ML analyzed the data and prepare the manuscript.

## Acknowledgements

Not applicable

## REFERENCES

1. Wahl B, O’Brien KL, Greenbaum A, Majumder A, Liu L, Chu Y, Luksic I, Nair H, McAllister DA, Campbell H, Rudan I, Black R, Knoll MD. Burden of Streptococcus pneumoniae and Haemophilus influenzae type b disease in children in the era of conjugate vaccines: global, regional, and national estimates for 2000-15. Lancet Global Health. 2018;6:e744–e757.

2. Oosterhuis-Kafeja F, Beutels P, Van Damme P. Immunogenicity, efficacy, safety and effectiveness of pneumococcal conjugate vaccines (1998-2006). Vaccine. 2007;25:2194–212.

3. Loo JD, Conklin L, Fleming-Dutra KE, Knoll MD, Park DE, Kirk J, Goldblatt D, O’Brien KL, Whitney CG. Systematic review of the indirect effect of pneumococcal conjugate vaccine dosing schedules on pneumococcal disease and colonization. Pediatric Infectious Diseases Journal. 2014;33:S161–71.

4. Bogaert D, De Groot R & Hermans PWM. *Streptococcus pneumoniae* colonisation: the key to pneumococcal disease. The Lancet Infectious Diseases. 2004;4:144–154.

5. Tsaban G, Ben-Shimol S. Indirect (herd) protection, following pneumococcal conjugated vaccines introduction: A systematic review of the literature. Vaccine. 2017;35:2882–2891.

6. Flasche S, Van Hoek AJ, Goldblatt D, Edmunds WJ, O’Brien KK, Scott JA, Miller E. The Potential for Reducing the Number of Pneumococcal Conjugate Vaccine Doses While Sustaining Herd Immunity in High-Income Countries. PloS Med. 2015;12:e1001839.

7. O’Brien KL When less is more: how many doses of PCV are enough? Lancet Infect. Dis. 2018;18:127–128.

8. Goldblatt D, Southern J, Andrews NJ, Burbidge P, Partington J, Roalfe L, Valente Pinto M, Thalasselis V, Plested E, Richardson H, Snape MD, Miller E. Pneumococcal conjugate vaccine 13 delivered as one primary and one booster dose (1 + 1) compared with two primary doses and a booster (2 + 1) in UK infants: a multicentre, parallel group randomised controlled trial. Lancet Infect. Dis. 2018;18:171–179.

9. Temple B, Toan NT, Uyen DY, Balloch A, Bright K, Cheung YB, Licciardi P, Nguyen CD, Phuong NTM, Satzke C, Smith-Vaughan H, Vu TQH, Huu TN, Mulholland EK. Evaluation of different infant vaccination schedules incorporating pneumococcal vaccination (The Vietnam Pneumococcal Project): protocol of a randomized controlled trial. BMJ Open. 2018;8:e019795.

10. Cobey, S et al. Niche and Neutral Effects of Acquired Immunity Permit Coexistence of Pneumococcal Serotypes. Science. 2012;335:1376–1380.

11. Lipsitch, M et al. No coexistence for free: Neutral null models for multistrain pathogens. Epidemics. 2009;1:2–13.

12. Croucher, N.J. et al. Population genomic datasets describing the post-vaccine evolutionary epidemiology of *Streptococcus pneumoniae*. Nature Scientific Data. 2015;2:150058.

13. Cai FY, Fussell T, Cobey SE, Lipsitch M. Use of individual-based model of pneumococcal carriage for planning a randomized trial of a vaccine. Submitted 2018.

14. Waight PA, Andrews NJ, Ladhani NJ, Sheppard CL, Slack MP, Miller E. Effect of the 13-valent pneumococcal conjugate vaccine on invasive pneumococcal disease in England and Wales 4 years after its introduction: an observational cohort study. Lancet Infect Dis. 2015;15:629.

15. Dagan, R et al. The effect of an alternative reduced-dose infant schedule and a second year catch-up schedule with 7-valent pneumococcal conjugate vaccine on pneumococcal carriage: A randomized controlled trial. Vaccine. 2012:30:5132-5140.

16. Rinta-Kokko H, Dagan R, Givon-Lavi N & Auranen K. Estimation of vaccine efficacy against acquisition of pneumococcal carriage. Vaccine. 2009;27:3831–3837.

17. Ota MO et al. The immunogenicity and impact on nasopharyngeal carriage of fewer doses of conjugate pneumococcal vaccine immunization schedule. Vaccine. 2011;29:2999–3007.

18. Cohen J. Statistical Power Analysis for the Behavioral Sciences. 2nd ed. Hillsdale, NJ: Lawrence Erlbaum Associates; 1988.

19. Search of: pneumococcal | (Map: Vietnam) - List Results - ClinicalTrials.gov. https://clinicaltrials.gov/ct2/results?cond=pneumococcal&map_cntry=VN. Accessed 14 July 2018.

20. Scott P, Herzog SA, Auranen K, Dagan R, Low N, Egger M, Heijne JC. Timing of bacterial carriage sampling in vaccine trials: a modelling study. Epidemics. 2014;9:8–17.

21. Auranen K, Rinta-Kokko H, Goldblatt D, Nohynek H, O’Brien KL, Satzke C, Simell, B, Tanskanen A, Käyhty H; Pneumococcal Carriage Group (PneumoCarr). Colonisation endpoints in Streptococcus pneumoniae vaccine trials. Vaccine. 2013;32:153–8.

22. Auranen K, Rinta-Kokko H, Goldblatt D, Nohynek H, O’Brien KL, Satzke C, Simell B, Tanskanen A, Käyhty H. Design questions for Streptococcus pneumoniae vaccine trials with a colonisation endpoint. Vaccine. 2013;32(1):159–64.

23. Mehtälä J, Antonio M, Kaltoft MS, O’Brien KL, Auranen K. Competition between Streptococcus pneumoniae strains: implications for vaccine-induced replacement in colonization and disease. Epidemiology. 2013;24:522–9.

24. Cai FY, Fussell T, Cobey SE, Lipsitch M. Use of an individual-based model of pneumococcal carriage for planning a randomized trial of a vaccine. PloS Computational Biology in press (2018).

25. Mehtala J, Antonio M, Kaltoft MS, O’Brien KL, Auranen K. Competition between Streptococcus pneumoniae strains: implications for vaccine-induced replacement in colonization and disease. Epidemiology. 2013;24:522–9.

26. Auranen K, Mehtala J, Tanskanen A, Kaltoft MS. Between-strain competition in acquisition and clearance of pneumococcal carriage—epidemiologic evidence from a longitudinal study of day-care children. Am J Epidemiol. 2010;171:169–76.

27. Lipsitch M. Interpreting results from trials of pneumococcal conjugate vaccines: a statistical test for detecting vaccine-induced increases in carriage of nonvaccine serotypes. Am J Epidemiol. 2001;154:85–92.

28. Halloran ME, Longini IM Jr, Struchiner CJ. Design and interpretation of vaccine field studies. Epidemiol Rev. 1992;21:73–88.

29. Halloran ME, Auranen K, Baird S, Basta NE, Bellan S, Brookmeyer R, Cooper B, DeGruttola V, Hughes J, Lessler J, Lofgren ET, Longini I, Onnela J, Ozler B, Seage G, Smith TA, Vespignani A, Vynnycky E, Lipsitch M. Simulations for Designing and Interpreting Intervetion Trials in Infectious Diseases. BMC Med. 2017;15:223.

30. Huang SS, Platt T, Rifas-Shiman SL, Pelton SI, Goldmann D, Finkelstein JA. Post-PCV7 changes in colonizing pneumococcal serotypes in 16 Massachusetts communities, 2001 and 2004. Pediatrics. 2006;116:e403–13.

